# Supercharged Fluorescent Protein-Apoferritin Cocrystals for Lighting Applications

**DOI:** 10.1101/2023.10.17.562704

**Authors:** Marta Patrian, Ahmed Shaukat, Mattia Nieddu, Jesús A. Banda-Vázquez, Jaakko V. I. Timonen, JP Fuenzalida-Werner, Eduardo Anaya□Plaza, Mauri A. Kostiainen, Rubén D. Costa

## Abstract

The design of lighting sources based on fluorescent proteins (FPs) has been limited by the lack of protocols to stabilize FPs under preparation (deposition techniques, organic solvents, *etc*.) and working (temperature, irradiation, *etc*.) conditions. As a critical bottleneck, photo-induced heat generation due to FP motion and quick heat transfer leads to working device temperatures of *ca*. 70 °C, resulting in a quick FP-denaturation and, in turn, a quick loss of the device performance. Herein, we showcase FP stabilization for lighting devices with an electrostatically self-assembled FP-apoferritin cocrystals embedded in a silicone-based color down-converting filter. This strategy highlights three major advances: *i*) engineering of positively supercharged FPs (+22) without losing photoluminescence and thermal stability compared to its native form, *ii*) a crystallization protocol resulting in highly emissive cocrystals keeping the photoluminescence features of the FPs, and *iii*) a 40-fold increase of the lighting device stability compared to reference devices due to the reduction of the device working temperatures to 40 °C. Thus, the success of this multidisciplinary approach contributes toward developing stable energy-related protein-based optoelectronics.

## Introduction

Though light-emitting diodes (LEDs) will dominate domestic and industrial artificial lighting, their eco-efficient development is a major concern. Major reasons are the lack of efficient recycling protocols and the need for rare-earth-based inorganic phosphors (IPs) as color down-converting filters.^1,2^ In this context, extensive research has focused on hybrid light-emitting diodes (HLEDs), in which organic phosphors based on small molecules,^3^ conjugated polymer,^4^ coordination complexes,^5,6^ *etc*. embedded into polymer and epoxy matrices have been explored. Among them, fluorescent proteins (FPs) are considered a paradigm of sustainable development with respect to their cheap bacterial production, easy recyclability, water-processability, and excellent emission merits: high molar extinction coefficients (ε) and photoluminescence quantum yields (□), as well as narrow emission band converting the whole visible spectrum.^7,8^ In addition, FP-based HLED performance stood up with stabilities of >3,000 h and efficiencies of >130 lm/W at low-power conditions^9^ (<50 mW/cm^2^ photon flux excitations), comparable to other devices with traditional OPs: *i*) perylene diimide-polymer with <700 h@130 lm/W, *ii*) BODIPYs-polymer with <10 h@13 lm/W, and *iii*) Iridium(III) complex-polymer with <1,000 h@100 lm/W.^3–6^ In contrast, the device stability is typically reduced to <5 min at high-power operation conditions (blue 200 mW/cm^2^ photon-flux excitation)^10^ due to photo-induced heat generation in the color down-converting coating related to FP motion and efficient heat transfer in a water-rich environment.^9,10^ Several strategies have been proposed to circumvent this issue by *i*) fabricating waterless polymer composites (<2 h)^9^ and *ii*) water-free FP isolation using sol-gel chemistry tools (<120 h).^11^

Herein, we explore the hypothesis of whether the immobilization of FPs within a protein cage crystal *via* electrostatic self-assembly could effectively restrict photo-induced heat generation towards enhanced high-power Bio-HLEDs. Protein cages are self-assembled nanocompartments that are used as ideal building blocks for crystalline assemblies due to their uniform size and shape.^12,13^ As a leading example, apoferritin (aFt) cages feature an inner cavity and crystal pores that can host a myriad of materials: *i*) ions and metals like Au, Zn, Ce or Ni or Co, ^14–18^ *ii*) organic molecules, such as photosensitizers, host-guest molecules, dendrimers^19–24^, and *iii*) biological building blocks, such as protein and peptides.^25–28^ In addition, cocrystallization is driven by electrostatic interactions, resulting in a modular and flexible concept for the formation of aFt-based assemblies with diverse chemical compositions and nanostructures.^29^ However, the cocrystallization with FPs has remained elusive up to date. Indeed, we have previously shown that supercharged fusion peptides (*i*.*e*., green fluorescent protein (GFP) fused with polylysine peptide) form ordered structures when combined with cowpea chlorotic mottle virus (CCMV).^30^ However, in the case of aFt cocrystals, the presence of GFP hindered the formation of crystalline assemblies, while attempts to engineer super positive GFP variants have led to outstanding thermal stabilities, but at the cost of the *ϕ*.^31^ This is in line with the design of supercharged enzymes that typically feature an enhanced thermostability, but a reduced activity and/or loss of functionalities.^32,33^

This work answers to these synergistic challenges by disclosing *i*) engineering approach for the design of a positively supercharged mGreenLantern (scmGL) variant featuring 22 positive charges without losing *ϕ*, and *ii*) a protocol to form aFt-scmGL cocrystals that keep the photoluminescence performance of the scmGL. Finally, aFt-scmGL cocrystals were easily implemented in a silicone-based color down-converting filter for high-power Bio-HLEDs, corroborating our initial hypothesis with a strongly reduced photo-induced heat generation from 65 °C to 40 °C working temperatures. This leads to 40-fold enhanced device stability compared to reference devices.

## Results and Discussion

### Positively supercharged mGreenLantern

Supercharging often requires extensive surface mutations on the proteins/enzymes, significantly impacting the equilibrium and extent of the non-covalent interactions that allow them to maintain their three-dimensional structure (fold).^31^ To overcome this issue, we first decided to determine the best supercharging starting point among archetypal enhanced green fluorescent protein (EGFP), super folder green fluorescent protein (sfGFP), and the recently published mClover variant called mGL (**Figure 1A**).^34–36^ To this end, we benchmarked them using the temperature of non-reversibility (T_nr_). The T_nr_ curves portray irreversible changes in protein structure, leading to changes in fluorescence after exposure to a specific temperature and subsequent cooling.^37^ As shown in **Figure 1B**, mGL is clearly superior to EGFP and sfGFP, both start losing their folding recovery capacity already before 50 °C. At the same time, mGL refolding capacity is only impaired above 80 °C. This results in an increased total area of T_nr_ (folding reversibility) going down from 61.8 (mGL), to 58.8 (sfGFP), and to 53.3 (EGFP). To further confirm the higher thermostability showcased by mGL, we measured the fluorescence overtime at 70 °C, mimicking the traditional Bio-HLED operating temperature (**Figure 1C**).^10^ Here, the superior folding stability of mGL became more prominent, showing a 23 % loss after 800 minutes compared to the 50 % and total loss for sfGFP and EGFP, respectively. Thus, we selected mGL as a starting point for supercharging.

**Figure 1.**
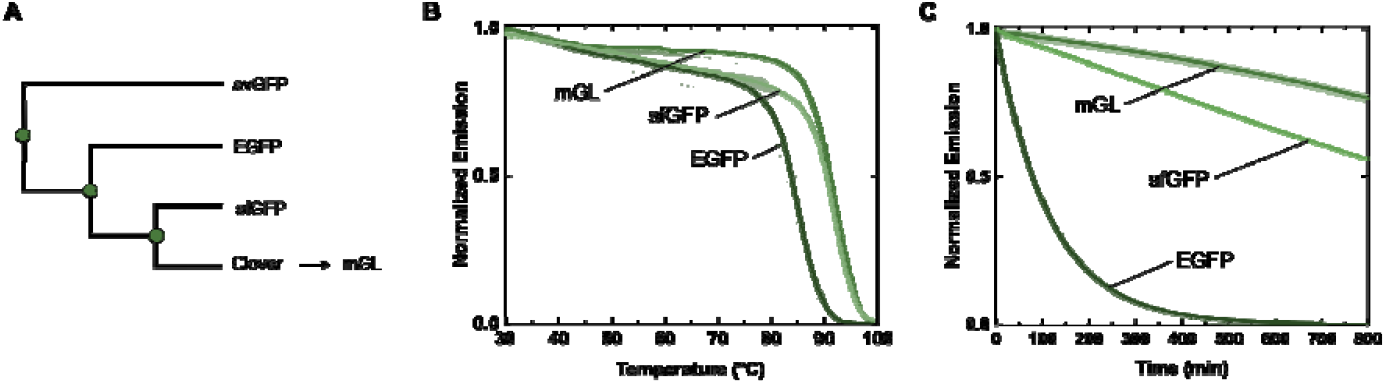
(A) Phylogenetic distance of aequorea victoria avGFP, EGFP, sfGFP, and mGL. (B) T_nr_ of mGL, sfGFP, and EGFP in PBS buffer solution. (C) Fluorescence overtime at 70 °C of mGL, sfGFP, and EGFP in PBS buffer solution.

The general approach to positively supercharge a protein is based on the identification of polar surface residues, solvent-exposed amino acids (*i*.*e*., D, E, N and Q), under the assumption that these positions will be able to accommodate a positively charged amino acid (*i*.*e*., R or K).^31^ A more advanced approach also considers the interaction between surface atoms to preserve the native folding.^38^ However, functionality losses can still be present. We hypothesize that, despite the general structure of a protein being preserved by conserving critical interaction between surface residues, small changes in the protein fold increase the free space around the chromophore, decreasing its stabilization and diminishing its brightness.^39,40^ For example, super-positive cerulean (+32) shows an opening in the barrel near the chromophore of 18.1 Å compared to 17.5 Å of the native protein. This opening correlates with increased free volume inside the barrel (**Figure S1**). To reduce this risk, we identified the amino acids near the chromophore based on their van der Waals radii to selectively replace only the polar amino acids that did not interact with any secondary structural elements containing the identified residues above, ensuring that the existing salt bridges remained unaffected (**Table S1**). Finally, we consider those mutations that are highly exposed to reduce interprotein interaction that might reduce long-range electrostatic interaction in the cocrystallization process. ^41^ All-in-all, our strategy leads us to design eleven mutations: eight on the flexible protein loops and three on two *β*-sheets (**Figure 2A-B**). This results in a theoretical protein charge of +22 without considering the terminal histidine tag (*i*.*e*., +28 in total). Rosetta minimization on AlphaFold models showed that the mutations have no adverse effect on the protein conformational stability in comparison to mGL (**Figure 2C**). In contrast, Cerulean and Cerulean +32 exhibited the conformation flexibility range almost double (**Figure S1**). More importantly, the *β*-barrel opening at the chromophore site remains constant at 17.5 Å, and our model shows that the free space around the chromophore did not increase after mutations (**Figure 2D**).

**Figure 2.**
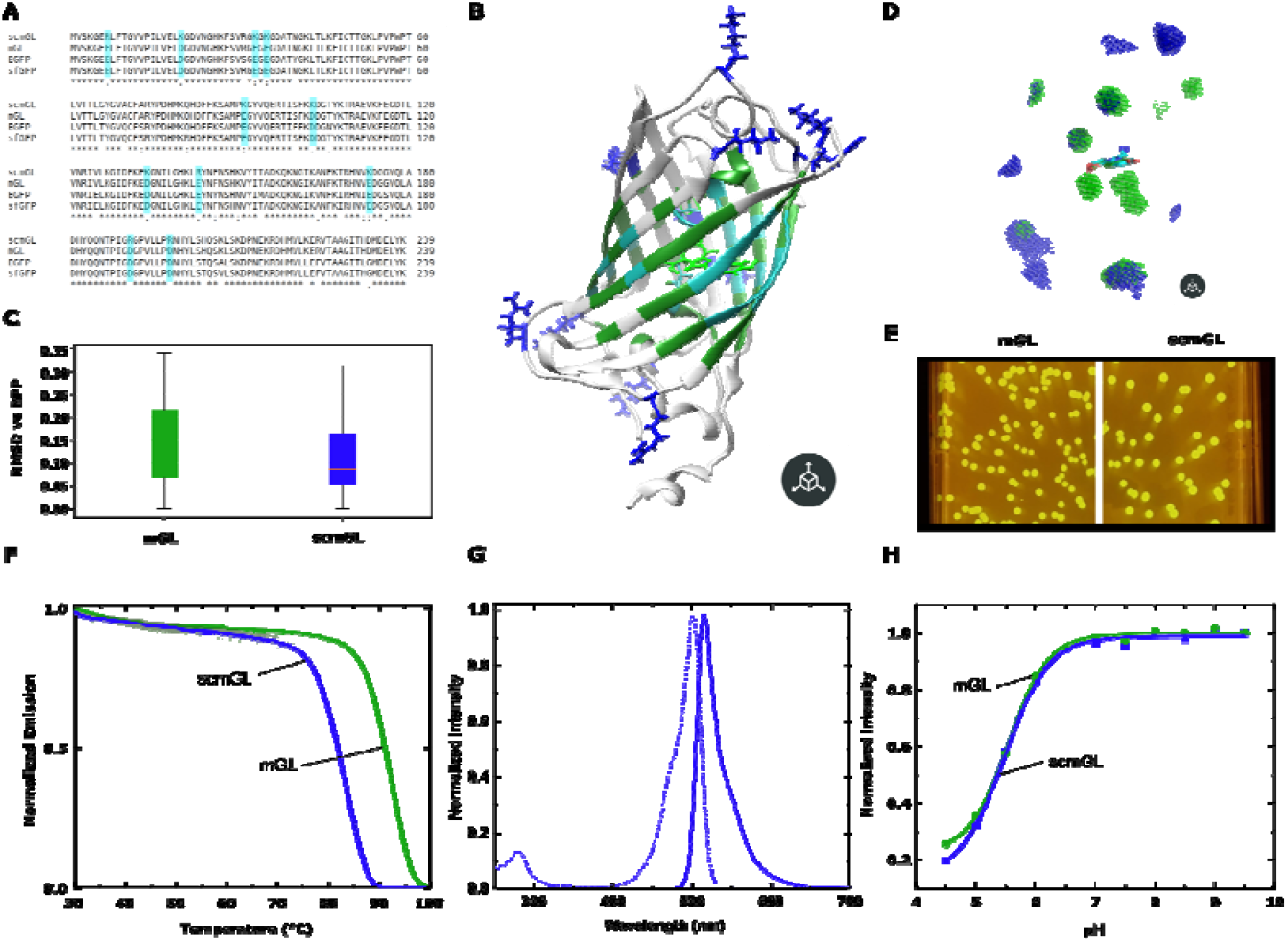
(A) Sequence alignment of mGL and scmGL. (B) Structural representation of scmGL, blue indicates the positively charged mutated amino acids, while dark green indicates the exclusion zone nearby the chromophore with light blue marking the amino acids in direct contact. (C) Root-mean-square deviation (RMSD) analysis of Rosetta minimized populations of mGL and scmGL. (D) Comparison of the internal and peripheral volume of mGL (green) and scmGL (scmGL). I *E. coli* colonies bearing mGL (left) and scmGL (right) expressing plasmids. (F) T_nr_ of mGL and scmGL. (G) Excitation and emission spectra of scmGL. (H) Fluorescence over the pH range 4.5–9.5 of mGL and scmGL.

Next, we cloned genes encoding mGL and scmGL to express them in *Escherichia coli*; both genes yielded intensely green fluorescent bacteria (**Figure 2E**). Following protein purification, the thermal and photophysical features of scmGL were measured using mGL as a reference. On one hand, the T_nr_ area of scmGL decreased from 61.8 to 52.8; the same level of EGFP (**Figure 2F**). The super-positive character most likely inhibits refolding in the absence of a prokaryotic environment. However, the mutations did not affect its capacity to withstand heat as observed by the similar T_m_ (65 °C (scmGL) *vs*. 66 °C (mGL); **Figure S2**). On the other hand, excitation and emission spectra strictly resemble the ones of the parental protein (**Figure 2G**). In addition, scmGL holds a high *ϕ* of 70 % as that of mGL (72%) along with a 5% lower *ε* (**Table 1**). Importantly, scmGL shows still higher *ϕ* and *ε* than sfGFP and EGFP with a brightness of 65.8 *vs*. 34.1 and 53.9, respectively. Finally, the pK_a_ (*i*.*e*., pH where only 50 % of its chromophore is in an emissive state) of scmGL does not show any variation (**Figure 2H** and **S3**), since all the extra ionizable groups are at least 20 Å away from the chromophore and primarily located in flexible loops. Consequently, scmGL conserves the broad active pH range of its parent mGL.

**Table 1.**
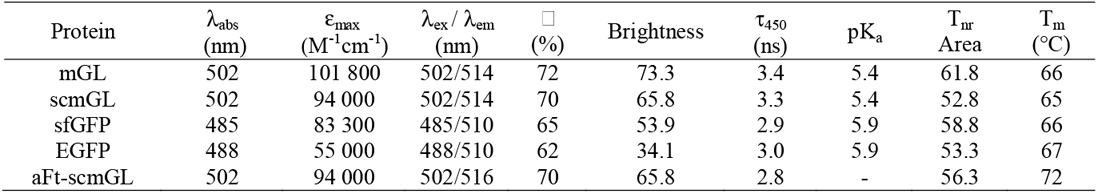
Overview of the photophysical and thermal figures of green FPs.

### aFt-scmGL cocrystallization

The initial investigation of the interaction between aFt protein cages and scmGL involved the application of dynamic light scattering (DLS). This technique enabled the monitoring of changes in scattering intensity (count rate) and the hydrodynamic diameter (*D*_h_), providing insights into the formation of significant complexes between the two entities. A constant concentration of aFt (0.1 mg□mL^-1^, 480 kDa, 0.20 µM) in Tris buffer (20 mM, pH 7.5) was titrated, increasing the scmGL concentration (**Figure 3B**). At first, aFt shows a rapid increase in particle count rate, which corresponds to the interaction of these two moieties and the creation of small aggregates. The scmGL amount required to complex aFt fully was 0.045 mg□mL^-1^ (1.5 µM). Based on the molecular weight of scmGL and aFt protein cages, there are *∼*7.5 scmGL proteins surrounding each aFt protein cage. To establish that the primary driving binding force is electrostatic interactions, the complexes were disassembled by gradually raising the ionic strength of the media, increasing the NaCl concentration (**Figure 3B**, right),^18^ reaching a total disassembly at 150 mM. The variation in the derived count rate is in good agreement with the change in *D*_h_ that increases with the scmGL concentration (**Figure 3C**, left). Upon increasing the scmGL concentration, the peak at 15.4 nm (native size of aFt) decreases, and a new peak at *∼*2 *μ*m (large aFt-scmGL complexes) evolves at 0.045 mg mL^-1^. At this point, the ionic strength was increased (**Figure 3C**, right), showing a decrease of *D*_h_ down to a similar size of free aFt (*i*.*e*., 13.4 nm at [NaCl] = 200 mM). As a control, the same experimental conditions were applied to prepare cocrystals of mGL and aFt without any success (Figure S4), as expected by the negative net charge of the mGL (*i*.*e*., -4). Thus, our scmGL design, in which the majority of the positive charges are located at the protein loops (**Figure 2B**), enables strong interaction through long-range electrostatic interactions driving the formation of the assembly.^41^

**Figure 3.**
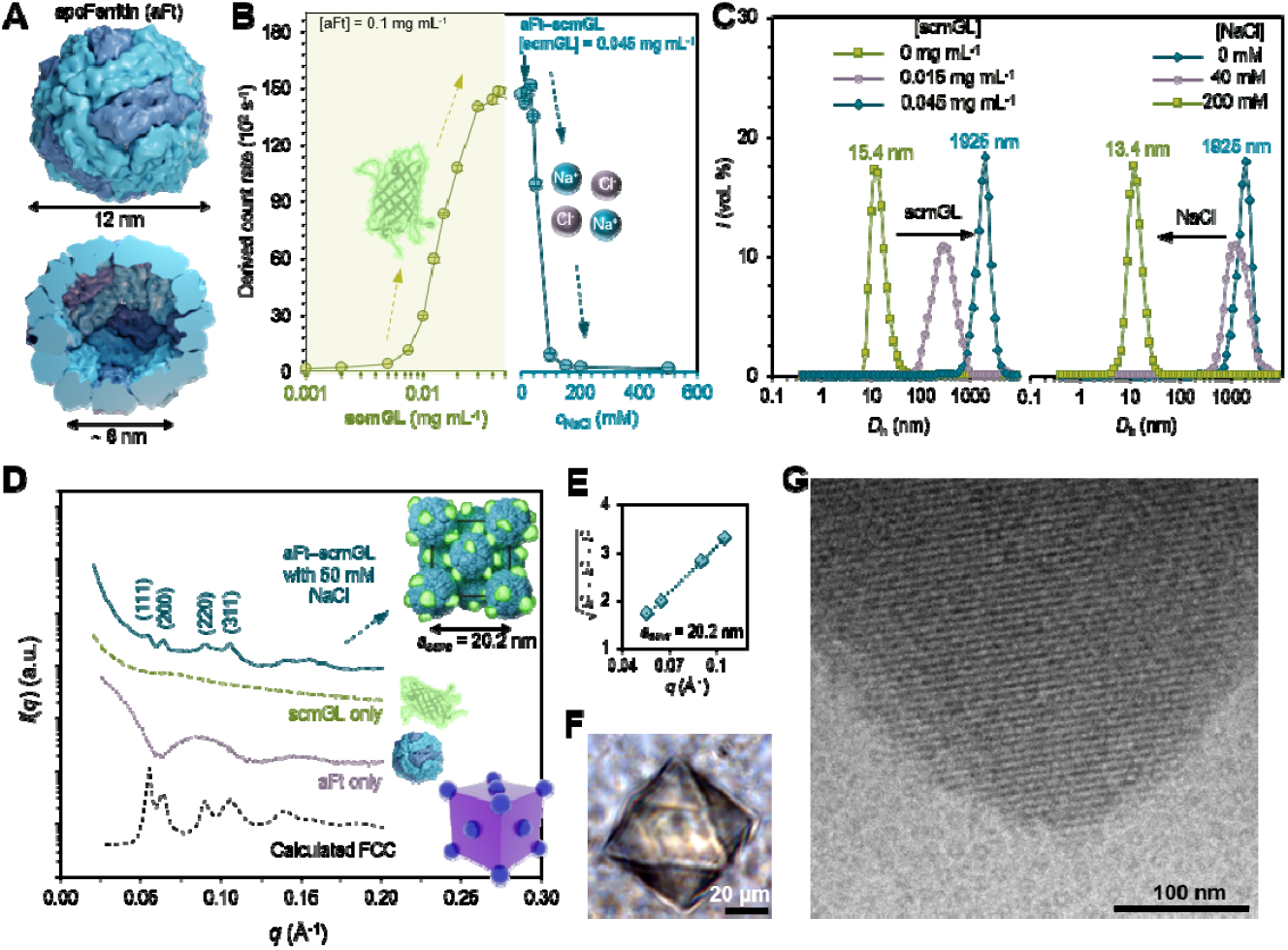
Self-assembly and structural characterization of aFt-scmGL. (A) Structure of the horse spleen apoferritin (aFt; PDB ID: 2W0O (B) Left: DLS of aFt solution (0.1 mg mL^-1^) titrated increasing scmGL concentration. Right: The aFt-scmGL are disassembled increasing the ionic strength of the medium with NaCl. (C) DLS data for the volume-averaged size distribution of free aFt titrated with an increasing amount of scmGL (left) and the resulting aFt-scmGL disassembled with NaCl (right). (D) SAXS diffractograms measured for aFt–scmGL complex at 50 mM of NaCl, compared to a FCC model, free aFt and scmGL (offset in the y-direction for clarity). Inset: FCC unit cell showing randomly distributed scmGL proteins attached to aFt protein cageI(E) Square root of the sum of the square of the Miller indexes of the assigned reflections for the FCC structure *vs*. the measured *q*-vector positions. Unit cell parameter of *a* = 20.2 nm and space group Fm3m no. 225. The concentrations in SAXS experiment is: aFt 4 mg□mL^-1^ and scmGL 1 mg□mL^-1^ in 20 mM Tris buffer (pH 7.5). (F) Optical microscopy and (G) Cryo-TEM image of vitrified aqueous solutions of aFt-scmGL complex at 50 mM of NaCl.

The crystalline structure of aFt-scmGL was determined by small-angle X-ray scattering (SAXS). Protein cages ([aFt] = 4 mg mL^-1^) were mixed with scmGL at 1 mg□mL^-1^ in Tris buffer (20 mM, pH 7.5) at varying NaCl concentrations to facilitate the formation of large assemblies. The SAXS patterns show clear Bragg reflections with well-resolved diffraction peaks at low NaCl concentrations ([NaCl] = 10–50 mM; **Figure S5**). In line with the above DLS assay, the SAXS pattern gradually faltered at higher NaCl as the assembly formation is hampered. The relative peak positions fit with the first allowed reflections of a face center cubic (FCC) lattice (**Figure 3D**). They are assigned to the (111), (200), (220), and (311) reflections, corresponding to ratios *q*_(hkl)_/*q*\* = √3, √4, √8, and √11, respectively. As shown **Figure 3E**, a lattice parameter determined for the FCC system is *a*_SAXS_ = 20.2 nm, while the calculated nearest-neighbor aFt center-to-center distance is 14.3 nm. This result is consistent with the diameter of aFt (∼ 12 nm) and the *D*_h_ obtained from the dynamic light scattering (DLS) analysis mentioned earlier. (15.4 nm; **Figure 3C**). Moreover, the confirmation of the Bravais lattice space group Fm3m (no. 225) is supported by comparing it to a simulated curve derived from a finite FCC structure. Thus, the FCC unit cell contains four aFt cages corresponding to an estimated amount of *ca*. 28 scmGL proteins per unit cell. Finally, the same octahedral crystal habit is noted for aFt–scmGL and aFt crystals (**Figures 3F**), suggesting that the [111] direction is thermodynamically favorable for the crystal growth. To further confirm the aFt arrangement, cryo-electron microscopy was employed. For aFt– scmGL sample ([NaCl] = 50 mM) a well-structured organization of individual aFt particles was noted (**Figure 3G**). To provide evidence of the homogenous presence of scmGL within the crystals, confocal microscopy was employed, acquiring multiple stacks of images around the focal plane (*i*.*e*., z-stack) of single aFt–scmGL crystals (**Figure 4A**). Here, the distinct planes (top, center, and bottom) illustrate the distribution of scmGL signal throughout the faceted octahedral crystal. Additionally, fluorescence microscopy analysis allowed us *i*) a direct comparison of the fluorescence signal between aFt and aFt-scmGL crystals (**Figure 4B**) and *ii*) a photobleaching experiment to study the stability of the crystal lattice (**Figure 4C**). On one hand, the fluorescence signal of aFt-scmGL crystals is 1000 times stronger than that of aFt crystals caused by the use of Cd^+2^ salts for the aFt crystallization.^42^ On the other hand, a defined crystal region was photobleached with a high-intensity laser beam (405 nm) without showing any recovery over time, indicating that the excess of unbound scmGL proteins in the surrounding environment did not diffuse and replace the photobleached scmGL proteins within the crystalline assembly. Thus, the aFt-scmGL crystal lattice is a static and stable structure.

**Figure 4.**
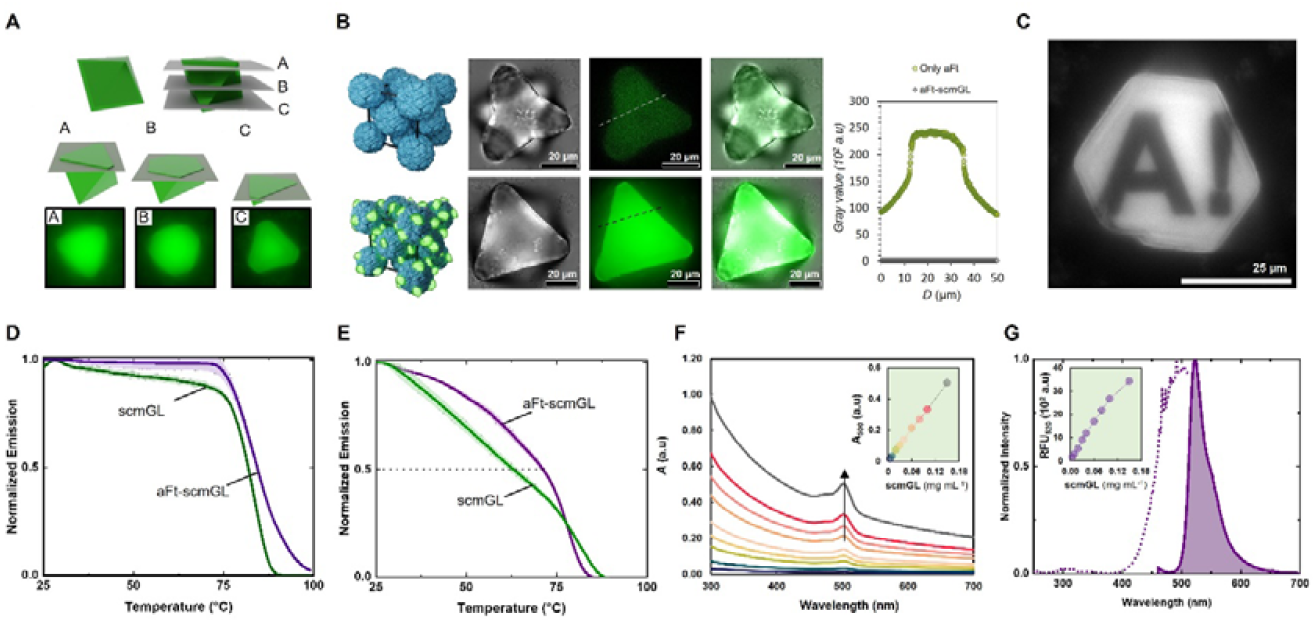
(A) Selected confocal microscopy z-stack planes of a single aFt-scmGL crystal showing that the scmGL is homogenously distributed in the crystal. (B) Comparison of aFt crystals and aFt-scmGL crystals. Top lane: bright-field, fluorescence, and composite images of the aFt crystals formed using CdSO_4_ [50 mM]. Bottom line: bright-field, fluorescence, and composite images of aFt-scmGL crystals with 50 mM NaCl. Right: Fluorescence intensity profiles (raw pixel values from the sCMOS camera) corresponding to the cross-sectional lines in the fluorescence images (center) of both aFt and aFt-scmGL crystals. (C) Fluorescence microscopy image of a selected area photobleached (Aalto University logo) on a the aFt-scmGL crystal, showing that the arrangement of the scmGL in the crystal is static and does not recover when photobleached. (D) T_nr_ of scmGL and aFt-scmGL cocrystals. (E) T_m_ of scmGL and aFt-scmGL cocrystals. (F) UV-Vis spectra of different concentrations of aFt-scmGL cocrystals assuming all scmGL protein is in the crystal. Inset: linear increase in 500 nm wavelength. (G) Excitation and emission spectra of aFt-scmGL cocrystals, Inset: linear increase in 520 nm emission wavelength with increase in cocrystals concentration assuming all scmGL protein is in the crystal.

Finally, we assess the thermodynamic and photophysical properties of aFt-scmGL crystals. Concerning the former, the aFt-scmGL shows an increase of the T_nr_ area from 52.8 to 56.3 and the T_m_ from 63 to 72°C, surpassing the T_m_ of mGL (**Figure 4D-E**). Thus, the protein packing aids the scmGL in retaining/recovering its folding at higher temperatures. The optical properties of the aFt-scmGL cocrystals were examined using steady-state and time-resolved UV-Vis absorption and emission spectroscopy. The UV-Vis absorption spectra (**Figure 4F**) display a linear increment of the absorption intensity at 500 nm upon adding more protein cocrystals, as noted for the unbound scmGL protein (**Figure S6**). Likewise, the aFt-scmGL features a well-structure emission spectrum similar to that of the parent scmGL protein in solution (**Figure 4G**) associated with the same *ϕ* at *ca*. 70 % (**Table 1**). As the only difference, the broadening of the excitation spectrum (**Figures 2 G** and **4G**) has been commonly attributed to FP aggregation phenomena that slightly affect the chromophore conformation.^9^ Indeed, the excited state lifetime (*τ*) is reduced due to the higher refractive index of the neighboring proteins in a multimeric aggregate (**Table 1**).^43^

### Implementation aFt–scmGL crystals in lighting devices

The most common color down-converting polymer matrix applied to Bio-HLEDs consists of a water-processed mixture of the stabilizer branched polyethylene oxide followed by the addition of a large molecular weight polyethylene oxide to control elastomeric and shape features. This mixture is stirred over a few hours and dried using a gentle vacuum procedure.^10^ To the naked eye, the aFt-scmGL crystals rapidly dissolve during the first stage. This could be related to synergistic effects: *i*) disruption of the polymer on the crystalline structure, *ii*) mechanical stress related to stirring, and *iii*) change of the local ionic strength due to the final dehydration processes that re-dissolve the crystals (*vide supra*). Capitalizing on the effective isolation of cocrystals in highly concentrated solutions, we use a biocompatible and transparent silicone matrix (ELASTOSIL®) applied in bio-imaging that features a reasonable water-compatibility (<20% v/v) without affecting optical features and a very mild curing process (*i*.*e*., room temperature and irradiation free) (**Figure 5A**). The success of this approach was confirmed by microscopic and photophysical assays (**Figure 5B-C** and **Table 2**). Confocal microscopy images confirmed the presence of crystals, in which the octahedral shape was mildly rounded (**Figure 5B**). This is attributed to shear forces during the mixing process and/or minor changes in the local ionic strength. Finally, the shape changes did not affect the fluorescence signal (**Figure 5C** and **Table 2**), keeping a green emission band associated to *ϕ* of 70 % and similar *τ* values to those noted in the cocrystals. However, the excitation spectrum exhibits a pronounced shoulder centered at around 395 nm that is ascribed to a fraction of protonated chromophore.^44,45^

**Table 2.** Overview of mGL, scmGL, and aFt-scmGL silicone-based filters and their respective Bio-HLEDs.

| Protein | $\lambda_{exc}/\lambda_{em}$<br>(nm) | □<br>(%) | $\tau_{450}$<br>(ns) | Device<br>stability<br>(min) | Device<br>temperature<br>(°C) |
| --- | --- | --- | --- | --- | --- |
| mGL | 502/527 | 67 | 3.3 | 1 | 65 |
| scmGL | 502/523 | 70 | 3.2 | 1 | 65 |
| aFt-scmGL | 510/530 | 69 | 2.7 | 40 | 40 |

**Figure 5.**
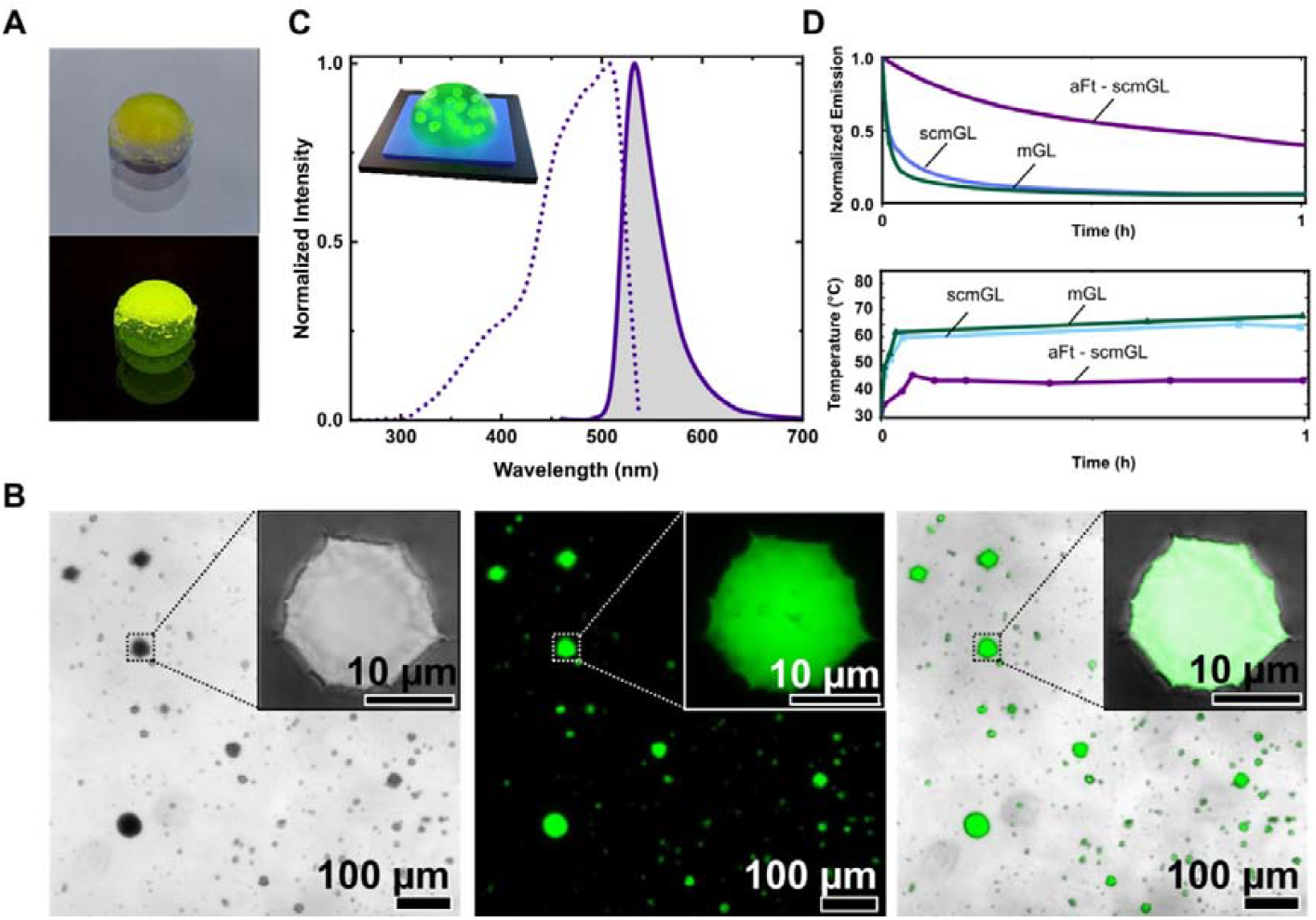
(A) Silicone-based coating with aFt-scmGL cocrystals under room illumination (top) and blue light (bottom). (B) Microscopy images of the aFt-scmGL filter with bright-field (left), confocal fluorescence (middle), and composite (right) images and their respective insets showing magnified image of one the crystals. (C) Excitation and emission spectra of silicone-based coating with aFt-scmGL cocrystals. (D) Device stability (top) and temperature (bottom) operating at high power conditions (200 mW/cm^2^).

Finally, green-emitting devices were fabricated using a blue-emitting LED chip (450 nm; Winger Electronics, 1W) that was directly covered by an optimized dome-shaped (6mm ⍰ and 1 mm; **Figure 5A**) silicone-processed aFt-scmGL (1 mg). At high-power driving conditions (200 mW/cm^2^ photon flux excitation), the device stability (*i*.*e*., 50 % loss of the initial intensity of the conversion band) was *ca*. 40 min in concert with a working temperature of around 40 °C (**Figure 5D**). This result is remarkable since reference devices with the same scmGL and mGL amounts showed stabilities of *<*1 min, reaching working temperatures of 65 °C (**Figure 5D**). The increase in device stability is related to the lack of photo-induced heat generation, as the crystal structure limits FP motion and heat transfer. Importantly, we analyzed the *post-mortem* coatings by microscopy and spectroscopic tools, encountering that *i*) the shape of the crystals holds with a detectable weak emission signal (**Figure S7**) and *ii*) a significant increase of the emission intensity at 450 nm related to the chromophore’s neutral form indicating that the primary deactivation mechanism was the photo-induced H-transfer process as noted in other FPs (**Figure S8**).^9,10^

## Conclusion

This work illustrates an innovative approach in enhancing protein-based lighting device performance with the design of *quasi* zero-thermal quenching color filters with cocrystals of aFt and super positively charged mGL as color down-converting materials. To realize this milestone, three essential advances have been achieved. At first, a new design concept for super positively charged FPs, in which peripheral amino acids are mutated, enables to preserve the chromophore environment, leading to a scmGL variant featuring +22 net charge and no loss in thermal and photoluminescence figures in contrast to the prior-art.^31^ In addition, these changes promote efficient long-range electrostatic interactions, resulting in the first cocrystal of aFt with FPs. What is more, these cocrystals feature a robust and static assembly with enhanced T_nr_ and T_m_ in concert with the same *ϕ* and emission band shape. This allow the fabrication of silicone-based color filters that were applied to on-chip high-power devices. They nicely outperformed the reference ones with a 40-fold enhanced stability due to the significant reduction of the working temperature from 65 ºC (FP filters) to 40 ºC (aFt-scmGL cocrystals). Overall, this work is seminal disclosing the first highly emissive all-protein-cocrystals and their earliest integration as active components in energy-related optoelectronics.

## Acknowledgment

This work has received funding from the European Research Council (ERC) under the European Union’s Horizon 2020 research and innovation programme (Grant Agreement No. 816856 (R.D.C) and No. 101002258 (MAK)). R.D.C. and J.B.V. acknowledge the European Union’s Horizon 2020 research for the MSCA grant AnBioLED No 101064305. M.N. and R.D.C. acknowledge FPNP-BioLED No. 101022975 funded by H2020-MSCA-IF-2020 of the European Commission. E.A.-P. acknowledge funding from Academy of Finland (Grant Agreement No. 341057). A.S, J.V.T., E.A.-P., and M.A.K. acknowledge the provision of facilities and technical support by Aalto University Bioeconomy Facilities and OtaNano-Nanomicroscopy Center (Aalto-NMC). We would like to acknowledge the Aalto University Internal Funding Call for Co-Operation initiatives with TUM.

## Author Contribution

M.P. A.S, and M.N. contributed equally; Project design: E.A.P, J.P.F-W, M.A.K. and R.D.C.; J.P.F-W. and M. P. conceptualized the genetic strategy, and planned and supervised experiments in protein engineering; M.P. conducted molecular biology, protein purification, and cell culture work, and helped with spectroscopy. J.A.B-V. performed computational analysis; M.A.K and E.A.-P. planned and supervised experiments in protein cage crystals and spectroscopy A.S. conducted co-crystallization and characterization experiments in protein cage crystals; J.V.T. and A.S. conducted experiments on microscopy. R.D.C. And M.N. planned and supervised experiments in material engineering. M.N. envisioned material design and performed device fabrication and characterization; Manuscript preparation: all authors; Funding acquisition: E.A.P, M.A.K, R.D.C., J.A.B-V, and M.N.

## Notes

### Competing Interest Statement

The authors have declared no competing interest.

