## Supplementary material for "Supercharged Fluorescent Protein-Apoferritin Cocrystals for Lighting Applications": SI_File

**Table of Content**

**Experimental section**

### Chemicals

Unless otherwise specified, all chemical reagents utilized in the experiments were obtained from Sigma-Aldrich and were used without any additional purification. The following were also obtained from Sigma-Aldrich: Tris base, horse spleen derived apoferritin (aFt) (received in 0.135 M NaCl solution, further diluted in milliQ water (aFT, 10 mg mL^-1^). Sodium chloride was obtained from VWR Chemicals. Milli-Q grade water was used in all experiments, while the buffer utilized in all experiments had a pH of 7.5 and consisted of 20 mM Tris base, which was adjusted using 1M HCl unless otherwise stated.

### Methods

#### FP production

All constructs used in this study were ordered as codon optimized synthetic genes (Twist Bioscience) cloned within the bacterial expression vector pET21(+). All the FPs were under the control of T7-LacO promoter and N-terminally tagged with 6x HisTag for subsequent protein purification. The FPs plasmid constructs were transformed in electro-competent Escherichia coli BL21 DE3 and plated on Luria Beltrani (LB) agar plates, containing 100 μg/ml Ampicillin. Single colonies were picked, inoculated in selective liquid LB medium, and grown shaking overnight at 30 °C. Subsequently, appropriate amounts of pre-culture were re-inoculated in fresh LB to an initial optical density (OD_600_) of 0.1 and grown at 30°C until OD_600_ 0.6 was reached. At this point, protein production was induced with 1 mM IPTG and the bacterial cells were incubated at 16 °C shaking for 48 hours.

#### Recombinant protein purification

Bacterial cells were harvested via centrifugation (25 minutes, 5000 g), and subsequently washed in PBS (NaCl 8 mg/L, KCl 0.2 g/L, Na_2_HPO_4_ 1.42 g/L, KH_2_PO_2_ 0.27 g/L, MQ water; pH=7.4). The pellets were stored at -20°C overnight before protein purification. Sonication was performed on ice at 80 Amplitude for a total process time of 8 minutes. After sonication, the disrupted cell suspension was centrifuged for 1 h at 18 000 rpm. His-tagged protein present in the supernatant were purified via HisTrap^TM^ (Äkta pure cytiva) and desalted via HiPrep^TM^ 26/10 Desalting column (Äkta pure cytiva). Following purification, fluorescent proteins were concentrated with Centrifugal Filter Units (Merck), frozen in liquid nitrogen, and stored at -80°C.

#### Thermocycler-based modulated scanning fluorimetry

Modulated Scanning Fluorimetry was performed as described in Svilenov *et al*.^1^ The Thermocycler CFX96 Touch Real-time PCR System (Bio-Rad) was employed to perform MSF measurements. One standard program composed of heating and cooling cycles ranging from 25 °C to 99 °C was used to measure the progressive loss of fluorescence and the irreversible unfolding of the FPs studied in this work. The samples were heated 5 °C/sec and held for 1 min at the temperature peak, followed by a recovery period of 5 min at 25 °C. Due to the high sensitivity of the Thermocycler detector and high QY of the FPs used in this study, only 1 μM of FPs were added per well to avoid saturation. The thermograms were buffer-subtracted and normalized by the highest fluorescence read-out of each sample. Data analysis was performed using Origin 2019 (OriginLab Corporation, Northampton, MA, USA). Mean values and standard deviations of quintuplicate were calculated and plotted. Melting curves were obtained plotting the fluorescence values obtained at peak temperatures, while non-reversibility curves were obtained plotting the fluorescence values obtained at 25°C. The nonreversibility temperatures (T_nr_) areas were determined via the integration tool available in the software.

#### Computational methods

RMSD comparisons: the CA atoms from the conserved glycines in fluorescent protein^2^ and the CA from the two flanking residues per chromophore were used for RMSD comparison of every relaxation against the best relaxed structure per protein population.

Cavity detection and Cavity Volume calculations: for Cerulean, Cerulean-32, mGL and scmGL protein structures, the ParKVFinder software^3^ was used with default values for whole protein exploration: whole protein mode, low resolution mode, prove in of 1.4 Å, probe out of 4.0 Å, volume cutoff of 5.0 Å^3^, and removal distance of 2.4 Å. The volumes in cavities of interest per protein were added for comparison purposes. ParKVFinder usage, along with cavity representations and structure superpositions, were carried out on PyMol 2.5.0.

#### Dynamic light scattering

The hydrodynamic diameter (*D*_h_) of the assemblies was measured using a Malvern Instruments DLS device (Zetasizer Nano ZS Series) with a 4 mW He-Ne gas laser at a wavelength of 633 nm and an avalanche photodiode detector at an angle of 173°. All experiments were carried at room temperature. PMMA cuvettes were used for the size measurements. Zetasizer software (Malvern Instruments) was used to obtain the particle size distributions: 0.1 mg mL^-1^ of aFt (with a final concentration of 0.25 mM NaCl) dissolved in buffer (20 mM Tris (pH 7.5)) was titrated with different mGL or scmGL concentrations (0.05, 0.1, 1, 10 mg mL^-1^) to reach the desired ratio (no dilution correction was done as the total addition did not exceed 5 % of sample volume), which was finally titrated with 5 M NaCl to disassemble the complex.

#### Sample preparation for cryo-TEM, SAXS, optical micrscopy, confocoal micrscopy and UV-Vis

For all the experiment involved complexing the FP and protein cages, the FP proteins was first dialyzed to 20 mM Tris pH 7.5 buffer.

The crystals were prepared by first mixing 1.5 μL of 0–800 mM NaCl solution and 1.5 μL of 10 mg mL^-1^ scmGL in a PCR tube and then adding 6 μL 20 mM Tris pH 7.5 buffer and then finally combining 6 μL of aqueous aFt solution (10 mg mL^-1^). The samples were gently mixed with a pipette. Immediately after mixing complexes are formed for 10–70 mM NaCl samples, while samples beyond 80 mM NaCl sample contained no visible complexes. The samples were then incubated in refrigerator for 24 hour to settle down the complexes and the sediments were used for further characterization. For Cryo-EM, in order to avoid overcrowding of sample on the TEM grid, the samples prepared from above mentioned procedure were further diluted three times using a 20 mM Tris pH 7.5 buffer. As a result, the final concentration of aFt in the samples was determined to be 1.33 mg mL^-1^.

#### Cryogenic transmission electron microscopy

The cryo-TEM images were collected using JEM 3200FSC field emission microscope (JEOL) operated at 300 kV in bright field mode with an Omega-type zero-loss energy filter. The images were acquired with Gatan Digital Micrograph software while the specimen temperature was maintained at -187 °C. The cryo-TEM samples were prepared by placing 3 μL aqueous dispersion of the sample on a 200-mesh Lacey carbon film on Copper TEM Grids (agar scientific) and plunge-frozen into liquid ethane using Leica grid plunger with 3 s blotting time under 100 % humidity. The grids with vitrified sample solution were maintained at liquid nitrogen temperature and then cryo-transferred to the microscope. The TEM grids were plasma cleaned (20 seconds oxygen plasma flash using a Gatan Solarus). Images were further processed using ImageJ software.

#### Small-angle X-ray scattering

The SAXS samples were measured using the Xenocs Xeuss 3.0 C device equipped with a GeniX 3D Cu microfocus source (wavelength λ = 1.542 Å) and EIGER2 R 1M hybrid pixel detector at a sample-to-detector distance of 0.6 m. One-dimensional SAXS data was obtained by azimuthally averaging the 2D scattering data and the magnitude of the scattering vector *q* is given by *q* = 4π sin*θ* / λ, where 2*θ* is the scattering angle. For all the measurements, the scattering vector *q* was calibrated using a silver behenate standard and the 2D scattering data were converted into SAXS curves by azimuthal averaging. The samples were sealed in 1 or 1.5 mm glass capillaries (Hilgenberg GmbH) that has limited scattering in the measured *q* region.

#### Optical microscopy

The Zeiss Axiovert A1 inverted microscope was used to perform imaging through optical microscopy. To prevent distortion of the crystal habit, a chamber-like area was created using double-sided tape on all four sides to hold the sample (3 µL) between the glass slide and coverslip.

#### Confocal fluorescence microscopy and photobleaching

The confocal fluorescence and brightfield imaging of the crystals was done using a spinning disk confocal microscope (Nikon Ti-E with Crest Optics X-Light V3 scanner and Hamamatsu Orca Flash 4.0LT camera) with photobleaching capabilities (Gataca iLas2) and 60x/1.2W objective lens. The system was controlled using Micro-Manager. Small volumes (3 uL) of the crystal samples (aFt and aFt-scmGL) in buffer were placed between two precision cover glasses (Thorlabs CG15KH) separated from each other using a double sided tape spacer and imaged under identical conditions. Fluorescence images were excited using 470 nm laser (LDI Laser Diode Illuminator) at 1% power level with exposure time of 5 ms. Z-stacks were collected similarly at 500 nm steps. The brightfield images were collected using the same microscope in the transmitted light mode using a red LED (Thorlabs). The selected area photobleaching was done using 405 nm laser (Coherent OBIS 405 nm LX 100mW) at 50% power by raster scanning the beam to form the desired area in ca. 500 ms defined in the software plugin (Gataca Modular).

The confocal fluorescence and brightfield imaging of the silicone based devices (fresh and post-mortem) was done using a point scanning laser confocal microscope (Zeiss LSM710 on Examiner frame). Fluorescence was excited using using 488 nm laser at 2.5% power and emission was collected above 493 nm with 2.5x/0.06 objective lens (low magnification images) and 40x/1.1W objective lens (high magnification images). Fresh sample was the silicone resin mixed with crystals cured on a glass coverslip. The post-morten sample was a bulk piece of moded resin containing crystals.

#### Preparation of FP-silicone phosphors

The FP-silicone filters were prepared by mixing 100 µL of ELASTOSIL® Part A and 20 µL of ELASTOSIL® Part B with 1 mg of aFt-scmGL and their respective mGL and scmGL in 20 μL. The mixture was placed in the desired mold dimensions and dried in air for three days to obtain a semi-spherical coating.

#### Photophysical characterization of solutions and coatings

The aFt-scmGL with 50 mM NaCl crystal samples were prepared as described above. The crystals were incubated in the refrigerator for 24 h for sedimenting the crystals. After the incubation, the supernatant was replaced with fresh buffer (20 mM Tris with pH 7.5). For measurements, the UV-Vis and fluorescence spectra were measured using a Cytation 3 plate reader (BioTek) using 96-well plates. Alternatively, absorption spectra were acquired with a UV-vis spectrometer UV-2600 (Shimadzu), using a wavelength range 200–800 nm, scan speed medium, threshold 0.01 and a slit width of 2.0. ε was determined by relative measurement. The photophysical studies were carried out using a FS5 Spectrofluorometer (Edinburgh Instruments) with the SC-10 module for solid samples, the SC-30 Integrating Sphere to determine ϕ, and the time-correlated single photon-counting or TCSPC (64.3 ps pulse width) module to determine *τ*. The data was then adjusted to a mono- or bi-exponential decay fit using Origin Software. To calculate the average lifetime for each FP-coating, the following equation was used${<\tau>}_{0}= \frac{\int_{0}^{x} t \sum a_{i} exp\left( -\frac{t}{\tau_{i}} \right)dt}{\int_{0}^{x} \sum a_{i} exp \left( -\frac{t}{\tau_{i}} \right)dt}= \frac{\sum a_{i}\tau_{i}^{2}}{\sum a_{i}\tau_{i}}$; where *a_i_* (λ) is the amplitude fractions and *τ _i_* are the lifetimes. The measurements were performed at room temperature.

#### Device characterization

The above protein-coatings were placed at zero distance from the 450 nm LED (Winger Electronics; 1W) or 590 nm LED (Winger Electronics; 1 W). The Bio-HLEDs were characterized using a Keithley 2400 as a current source, while the changes in the electroluminescence spectrum were monitored using an AVS-DESKTOP-USB2 (Avantes) or Avantes Spectrometer 2048L (300 VA grating, 200 µm slit, CCD detector) in conjunction with a calibrated integrating sphere Avasphere 30-Irrad. The changes in the aFt-scmGL-based coating temperature were monitored using a thermographic camera T430sc (FLIR) coupled to the measuring system. Up to four replicates were measured for each configuration to exclude external influences.

### Supplementary Information – Figure S1

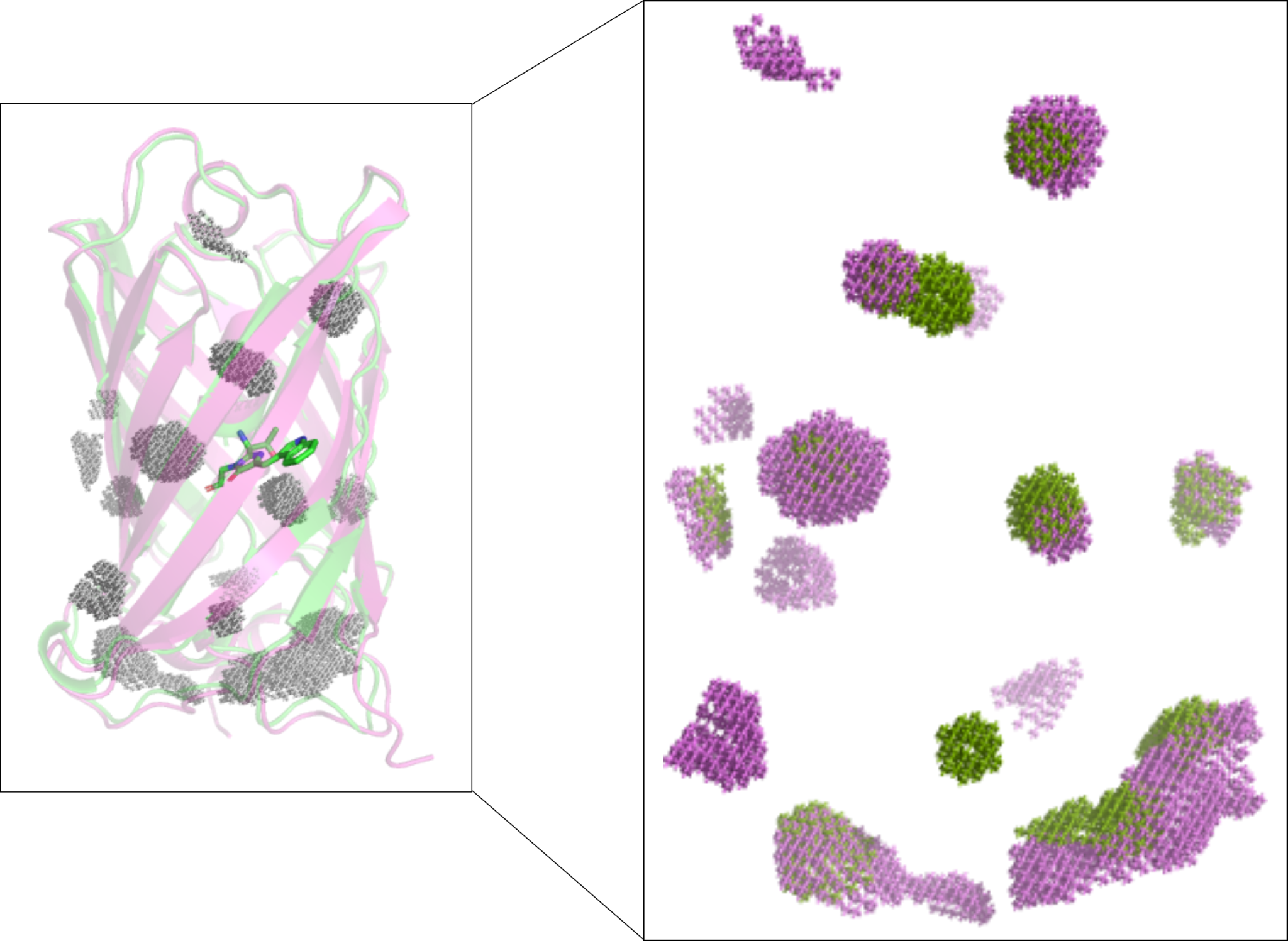

**Figure S1.** Cerulean Volume Comparison. Overlay structure super positive fluorescent protein cerulean in violet (PDB: 6MDR:A, net charge +32) and WT Cerulean in green (PDB: 2WSO). Zoom in on the available free volume in the β-barrel structure with the same color code. WT cerulean has significantly less volume available (left), indicating a more compacted structure than the superpositive counterpart.

### Supplementary Information – Table S1

**Table S1.** Comparison between H-bonds present on mGL (upper table) and scmGL (bottom table).

| **Donor** | **Acceptor** | **D-A distance (A)** |
| --- | --- | --- |
| LYS 79.A NZ | ASP 76.A OD1 | 3.240 |
| LYS 85.A NZ | ASP 82.A OD2 | 2.711 |
| ARG 109.A NH1 | GLU 111.A OE1 | 3.085 |
| ARG 122.A NH2 | GLU 115.A OE2 | 2.450 |
| LYS 126.A NZ | ASP 21.A OD2 | 2.972 |
| LYS 166.A NZ | ASP 180.A OD1 | 2.309 |
| LYS 166.A NZ | ASP 180.A OD2 | 2.881 |
| ARG 215.A NE | GLU 213.A OE1 | 2.945 |
| ARG 215.A NH2 | GLU 213.A OE2 | 3.157 |

| **Donor** | **Acceptor** | **D-A distance (A)** |
| --- | --- | --- |
| LYS 79.A NZ | ASP 76.A OD1 | 3.189 |
| LYS 85.A NZ | ASP 82.A OD2 | 2.794 |
| ARG 109.A NH1 | GLU 111.A OE1 | 3.118 |
| ARG 122.A NH2 | GLU 115.A OE2 | 2.433 |
| LYS 126.A NZ | ASP 21.A OD2 | 2.915 |
| LYS 166.A NZ | ASP 180.A OD1 | 2.380 |
| LYS 166.A NZ | ASP 180.A OD2 | 2.954 |
| ARG 215.A NE | GLU 213.A OE1 | 2.925 |
| ARG 215.A NH2 | GLU 213.A OE2 | 3.125 |

### Supplementary Information – Figure S2

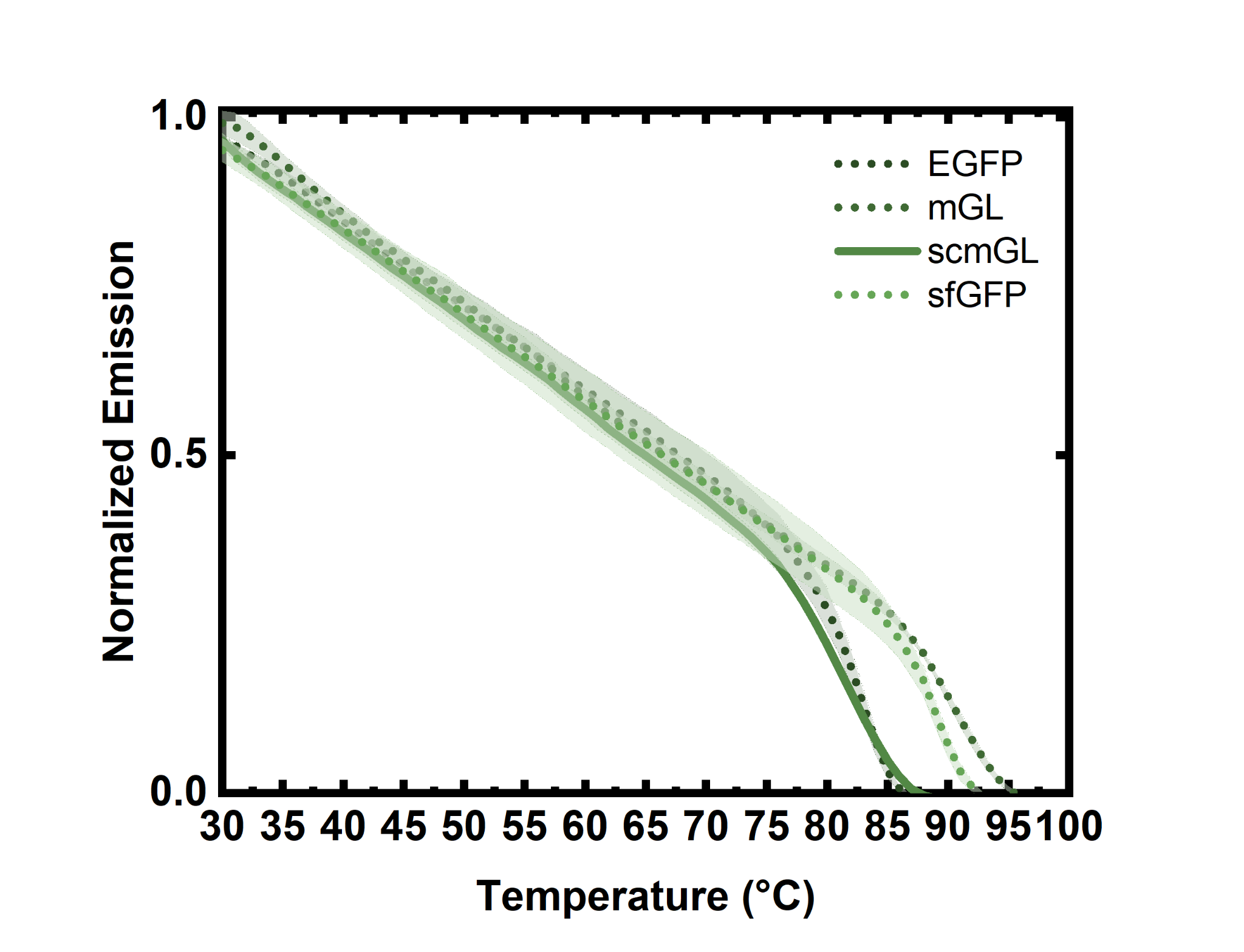

**Figure S2.** MSF of EGFP, mGL, sfGFP, and scmGL, monitoring the fluorescence intensity upon increasing temperature. T_m_ is calculated at 50 % of the emission intensity loss.

### Supplementary Information – Figure S3

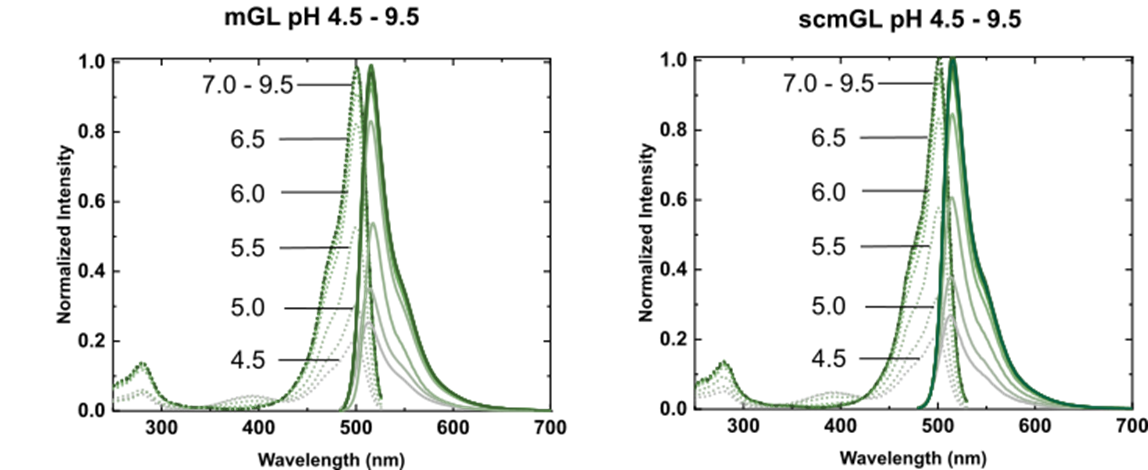

**Figure S3.** Excitation and emission spectra of mGL (left) and scmGL(right) measured in the pH range from 4.5 to 9.5.

### Supplementary Information – Figure S4

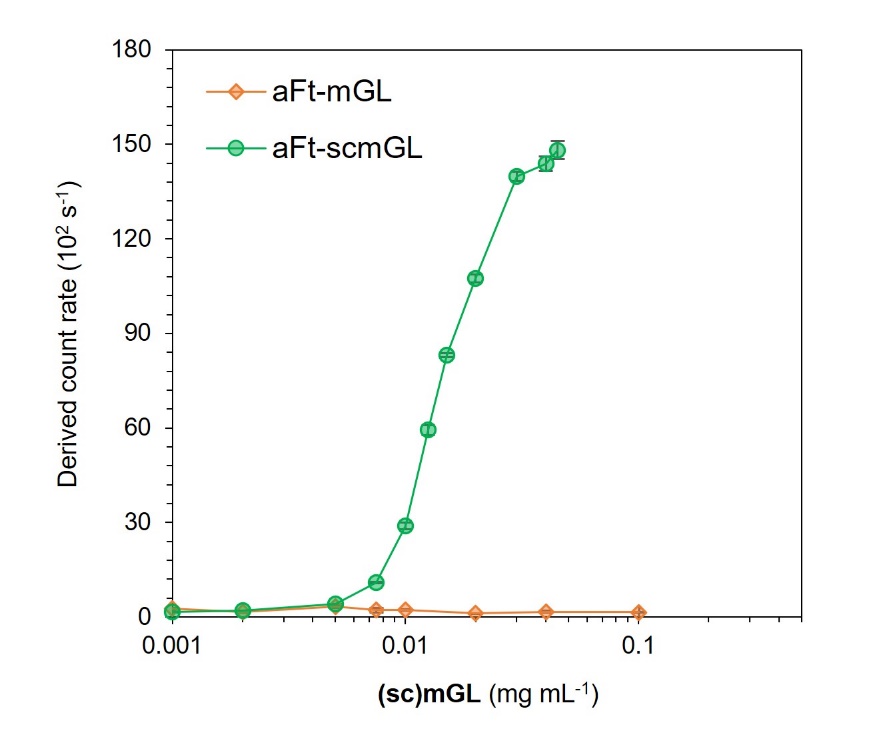

**Figure S4.** DLS showing the interaction of aFt with mGL with (scmGL) and without (mGL) charged groups.

### Supplementary Information – Figure S5

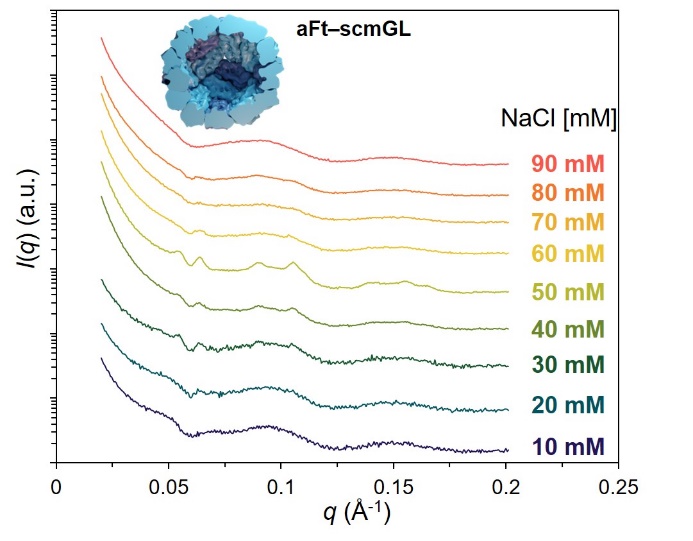

**Figure S5.** SAXS diffractograms of aFt-scmGL changing NaCl concentration.

### Supplementary Information – Figure S6

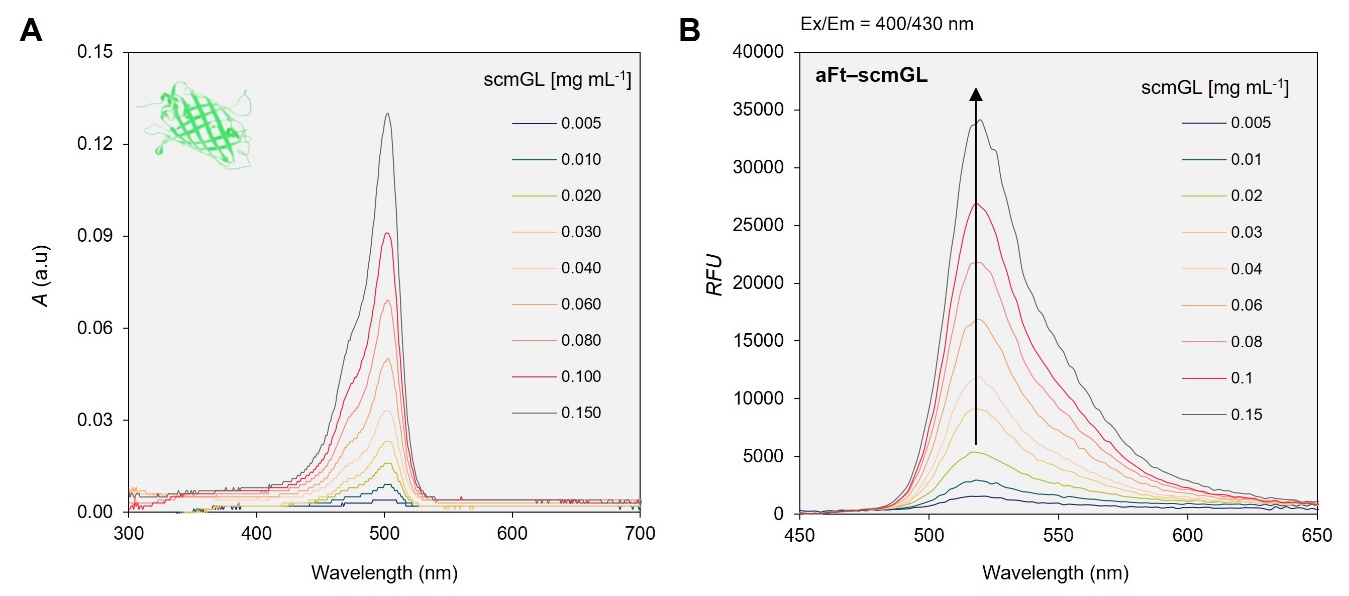

**Figure S6.** Optical propertise of scmGL UV-Vis absorption spectra of scmGL protein at various concentration ranging from 0.005 to 0.150 mg mL^-1^. . B) Fluorescence emission spectra of the same amount of scmGL in crystals. The sample was excited at 400 nm and the emission spectrum was collected from 430 nm

### Supplementary Information – Figure S7

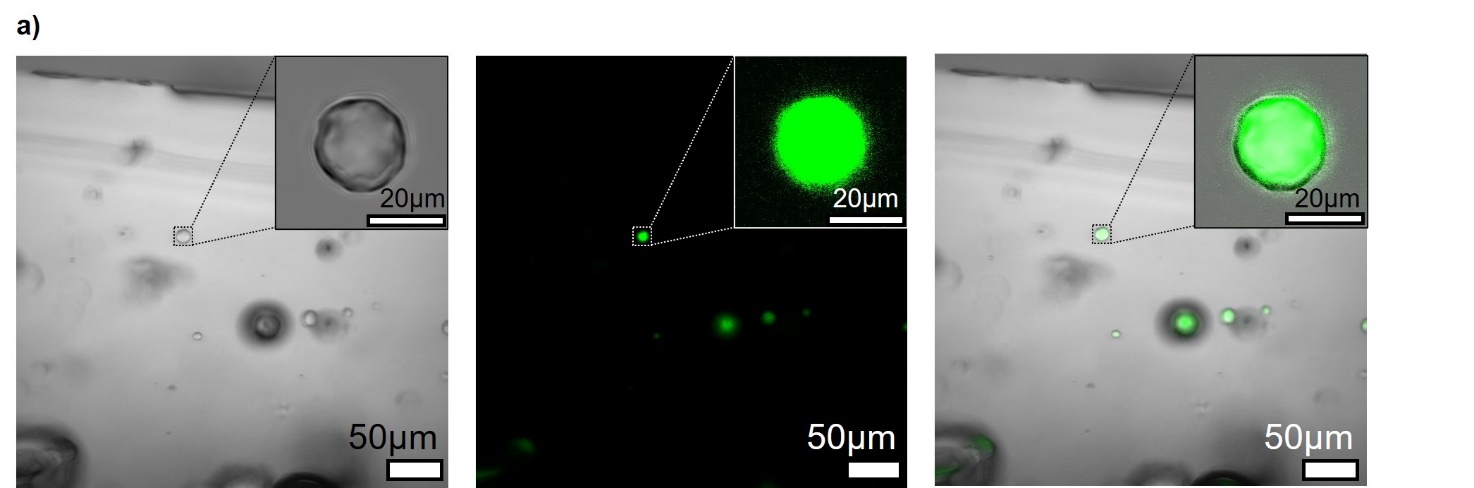

**Figure S7.** Microscopy image of a *post-mortem* aFt-scmGL-silicone filter with the bright-field (left), confocal fluorescence (middle), and composite (right) images and their respective insets showing magnified image of one of the small crystals.

### Supplementary Information – Figure S8

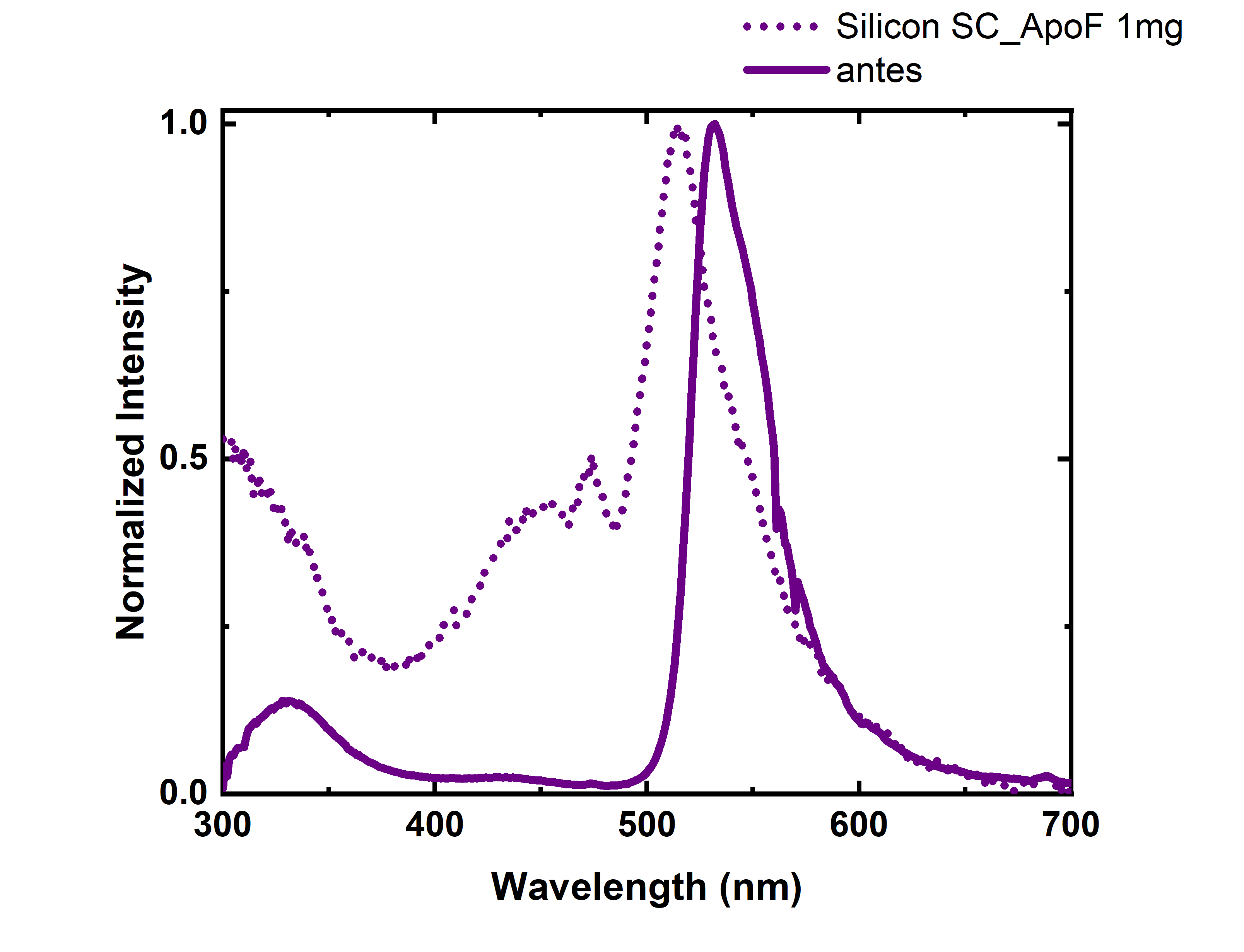

**Figure S8.** Emission spectra of fresh (solid line) and *post-mortem* (dotted line) aFt-scmGL silicone-based coatings at 280 nm excitation.
